# Interference with lipoprotein maturation sensitizes methicillin-resistant *Staphylococcus aureus* to human group IIA secreted phospholipase A_2_ and daptomycin

**DOI:** 10.1101/2022.02.06.476181

**Authors:** Marieke M. Kuijk, Yongzheng Wu, Vincent P. van Hensbergen, Gizem Shanlitourk, Christine Payré, Gérard Lambeau, Jennifer Herrmann, Rolf Müller, Jos A.G. van Strijp, Yvonne Pannekoek, Lhousseine Touqui, Nina M. van Sorge

## Abstract

Methicillin-resistant *Staphylococcus aureus* (MRSA) has been classified as a high priority pathogen by the World Health Organization underlining the high demand for new therapeutics to treat infections. Human group IIA secreted phospholipase A_2_ (hGIIA) is among the most potent bactericidal proteins against Gram-positive bacteria, including *S. aureus*. To determine hGIIA-resistance mechanisms of MRSA we screened the Nebraska Transposon Mutant Library using a sublethal concentration of recombinant hGIIA. We identified and confirmed the role of *lspA*, encoding the lipoprotein signal peptidase LspA, as a new hGIIA resistance gene in both *in vitro* assays and an infection model in hGIIA-transgenic mice. Increased susceptibility of the *lspA* mutant was associated with faster and increased cell wall penetration of hGIIA. Moreover, *lspA* deletion also increased susceptibility to daptomycin, a last-resort antibiotic to treat MRSA infections. Exposure of MRSA wild-type to the LspA-specific inhibitors globomycin and myxovirescin A1 induced a *lspA* mutant phenotype with regard to hGIIA and daptomycin killing. Analysis of >26,000 *S. aureus* genomes showed that LspA is highly sequence-conserved, suggesting that LspA inhibition could be applied universally. The role of LspA in hGIIA resistance was not restricted to MRSA since *Streptococcus mutans* and *Enterococcus faecalis* were also more hGIIA-susceptible after *lspA* deletion or LspA inhibition, respectively. Overall, our data suggest that pharmacological blocking of LspA may disarm Gram-positive pathogens, including MRSA, to enhance clearance by innate host defense molecules and clinically-applied antibiotics.

## Introduction

Infectious diseases are a significant cause of morbidity and mortality worldwide and are estimated to increase tremendously in the coming decades due to the rise of antimicrobial resistance [1]. The rapid development of antibiotic resistance does not just limit the success of treatment but also of prophylaxis of infections. Methicillin-resistant *Staphylococcus aureus* (MRSA) is a prominent example of a bacterium that has developed rapid antibiotic resistance over the past decades [2, 3]. Indeed, MRSA is ranked as one of the high priority pathogens by the World Health Organization with regard to the need for new therapeutic strategies [4]. While this bacterium is a common member of the human microbiota and asymptomatically colonizes the skin, gut, and nasal cavity, it can cause a wide spectrum of clinical diseases both in the hospital and in the community once *S. aureus* breaches host barriers.

The discovery of new antibiotics is slower than the emergence of new resistance mechanisms of pathogens [5-7]. Antibiotics are classified as substances that are able to kill bacteria (bactericidal) or inhibit their growth (bacteriostatic) [5]. Consequently, antibiotics target molecules or processes in the cell that are either essential or at least critical for the growth of bacteria. An alternative strategy to target bacterial pathogens could include anti-virulence or sensitizing drugs. These drugs may not affect bacterial viability or growth under laboratory conditions, but would affect bacterial fitness or even allow killing of bacteria in the context of specific host immune components, thereby clearing the infection. Indeed, *S. aureus* expresses a wide array of virulence molecules allowing for persistence in different host compartments through interference with a range of immune defense mechanisms and molecules [8].

The human group IIA secreted phospholipase A2 (hGIIA, also known as sPLA_2_-IIA) is a bactericidal enzyme that represents an important innate host defense molecule [9, 10]. hGIIA is highly cationic and effectively kills Gram-positive bacteria through hydrolysis of bacterial membrane phospholipids [11]. The enzyme is constitutively present at low levels (<5 ng/mL) in the blood circulation and its concentration increases rapidly to levels as high as 1000 ng/mL upon bacterial infection associated with sepsis [12, 13]. hGIIA requires anionic structures in the bacterial cell wall for binding to and penetration of the Gram-positive cell wall [14, 15]. Once at the membrane, hGIIA hydrolyzes membrane phospholipids resulting in bacterial lysis. hGIIA has been implicated in host defense against *S. aureus*. First, blocking hGIIA in acute phase serum results in loss of bactericidal effects against *S. aureus*, whereas addition of hGIIA to normal serum conferred anti-staphylococcal activity [16]. A bactericidal role of hGIIA has also been observed at barrier sites for example in human tears [17]. Second, hGIIA-transgenic (Tg) mice show higher survival rates compared to control littermates that are naturally sPLA_2_-IIA-deficient, after an experimental lethal dose of *S. aureus* [18, 19]. As a result, *S. aureus* has evolved resistance strategies against hGIIA-mediated killing, which are geared towards changing the overall charge of the membrane or cell wall. For example, *S. aureus* increases its surface charge by adding D-alanine residues to teichoic acids through the DltABCD machinery and L-lysine residues to membrane phospholipids through the activities of the enzyme MprF [14, 20]. The two-component regulatory system GraRS controls the expression of both *mrpF* and *dltABCD*, thereby controlling *S. aureus* resistance to cationic antimicrobial peptides and proteins such as hGIIA [21, 22]. Interestingly, the same bacterial genes are involved in *S. aureus* resistance to daptomycin, the antibiotic of last-resort to treat MRSA infections. Indeed, increased expression or gain-of-function mutations in *mprF* and *dltABCD* confer daptomycin non-susceptibility to *S. aureus* [23, 24]. Therefore, insight into hGIIA resistance mechanisms could provide new clues for the resistance against clinically-important antibiotics.

*S. aureus* is predicted to express between 50 to 70 lipoproteins, many of unknown function [25, 26]. Some lipoproteins are involved in antibiotic resistance, for example the beta-lactamase BlaZ and Dsp1 [27-29]. Before lipoproteins are considered mature, they need to be sequentially processed by the prolipoprotein diacylglyceryl transferase (Lgt) and lipoprotein signal peptidase II (LspA) enzymes. Lgt anchors prolipoproteins into the cell membrane through diacylglycerol and LspA subsequently generates the mature lipoprotein by removal of the signal peptide [30]. Both enzymes are conserved in all bacteria and marked as essential in Gram-negative but not Gram-positive bacteria [30]. Nonetheless, incorrect processing of lipoproteins changes the immune interaction of *S. aureus*; the deletion of *lgt* results in hypervirulence, whereas mutation of *lspA* attenuates virulence in a murine systemic infection model [31]. In addition, two screens, one designed to identify virulence genes and the other to identify MRSA resistance mechanisms to polymyxin B-mediated killing, identified *lspA* as a resistance determinant [32, 33].

The mechanisms by which *S. aureus* or MRSA resist hGIIA-mediated killing have never been studied in a comprehensive unbiased manner. Here, we screened the Nebraska Transposon Mutant Library (NTML) to identify hGIIA-susceptible mutants [34]. In addition to previously implicated genes [14, 35], we identified and confirmed that deletion of *lspA*, which we show to be extremely sequence-conserved, sensitizes *S. aureus* to hGIIA-mediated killing both *in vitro* and *in vivo*. Moreover, LspA confers resistance to the last-resort antibiotic daptomycin. Both hGIIA- and daptomycin susceptibility could be induced by treatment of MRSA with the LspA inhibitors globomycin and myxovirescin A1. The contribution of LspA to hGIIA resistance was not *S. aureus*-specific but was also observed in *Streptococcus mutans* (*S. mutans*) and in *Enterococcus faecalis* (*E. faecalis*). In conclusion, we identify LspA as a possible new therapeutic target to break resistance of *S. aureus* and possibly other Gram-positive pathogens to both endogenous antimicrobials and antibiotics routinely used in clinic.

## Materials and Methods

### Materials

Recombinant hGIIA was produced as described previously [36]. HEPES and CaCl_2_ were purchased from Sigma Aldrich and Merck, respectively. Albumin Bovine Fraction V, pH 7.0 (BSA) was purchased from Serva. SYTOX Nucleic acid stain was purchased from ThermoFisher and DiOC_2_(3) was obtained at Promokine / Bio-Connect B.V.. All antibiotics (chloramphenicol, erythromycin, daptomycin, gentamicin, and globomycin) were purchased from Sigma Aldrich.

### Bacterial culture

The NTML [34] was grown in Tryptic Soy Broth (TSB, Oxoid) supplemented with 5 μg/mL erythromycin. All other *S. aureus* strains and *Enterococcus* species (*E. faecalis* and *E. faecium*) used in this study (Table 1) were grown in Todd-Hewitt Broth (THB, Oxoid) with continuous shaking at 37°C. After overnight culture, strains were sub-cultured to an optical density at 600 nm (OD_600_) of 0.4 (early logarithmic phase; ≈1×10^8^ colony-forming units (CFU)/mL). The plasmid complemented strains were grown in THB supplemented with 20 μg/mL chloramphenicol. *S. mutans* was grown statically in Brain Heart Infusion (BHI) at 37 °C with 5% CO_2_. The following day, sub-cultures were grown to OD_600_ of 0.2 (early logarithmic phase). Plasmid complemented *S. mutans* strains were grown in the presence of 3 μg/mL chloramphenicol. *Escherichia coli* (*E. coli*) strains were grown in Lysogeny broth (LB) medium supplemented with appropriate antibiotics with continuous shaking.

**Table 1.**
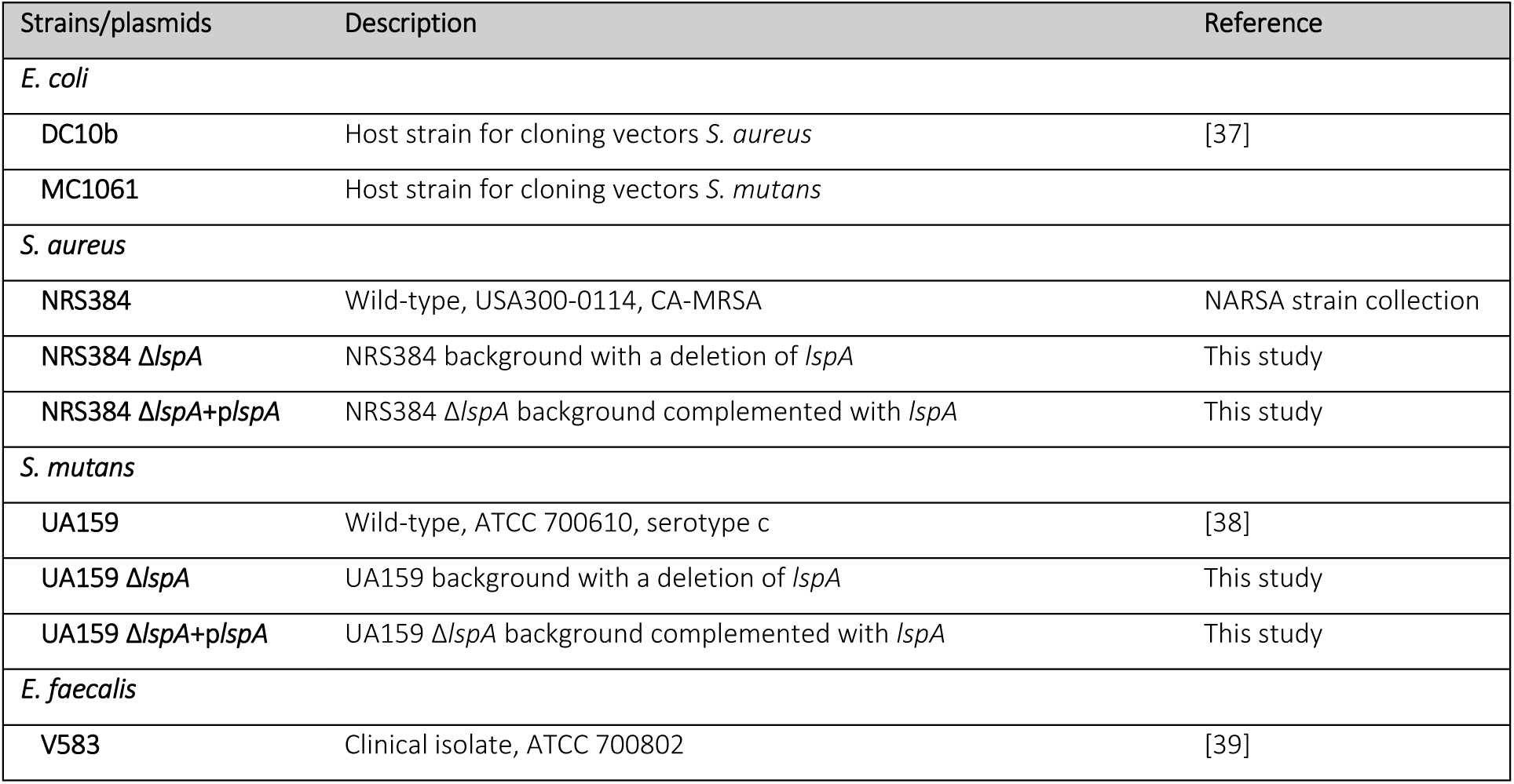
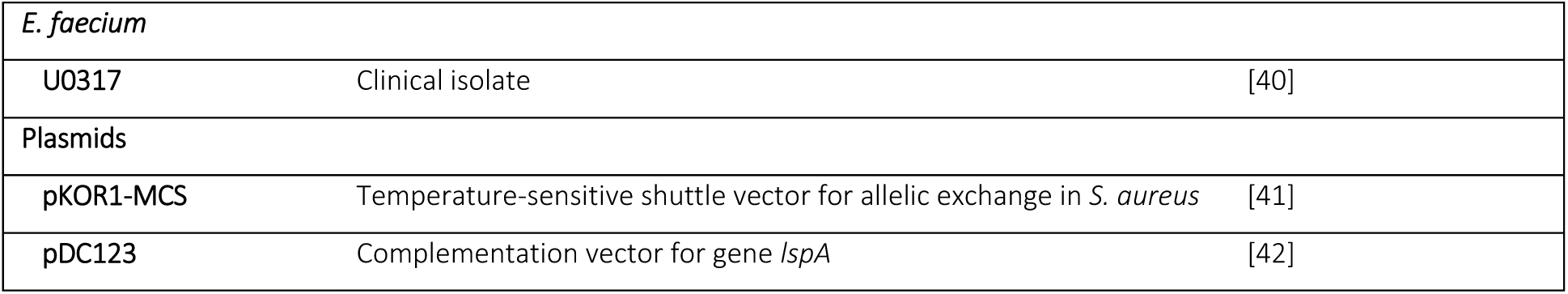
Overview of strains and plasmids used in this study.

### Screening the NTML for MRSA hGIIA resistance genes

All 1,920 mutants of the NTML were grown overnight in 96-well round bottom plates. After overnight culture, all transposon-mutant cultures were diluted 20 times in TSB supplemented with 5 μg/mL erythromycin and grown to early exponential phase. Cultures were subsequently diluted 20-fold in HEPES solution (20 mM HEPES, 2 mM CaCl_2_, pH=7.4) and exposed to 1.25 μg/mL recombinant hGIIA. After incubation for 1 hour at 37 °C, 5 μL droplets were plated on TS agar plates. Mutants with visibly reduced number of CFU were identified as putative hGIIA sensitive mutants.

### Construction of *lspA* deletion and *lspA* complemented strains

The markerless *lspA* (SAUSA300_1089) deletion mutant (MRSA Δ*lspA*) was generated in *S. aureus* strain USA300 NRS384. The temperature-sensitive and modified pKOR1 plasmid was used as described earlier [41, 43]. A fusion PCR of the upstream region of 1,008 base pairs (bp) and downstream region of 986 bp flanking the *lspA* gene was generated using NRS384 genomic DNA as template. The fusion PCR product was ligated into the pKOR1-MCS plasmid and amplified in *E. coli* DC10b before electroporation into *S. aureus* NRS384. Allelic exchange was performed through temperature shifts and counter selection [43].

To generate a *lspA* (SMU_853) deletion mutant in *S. mutans* strain UA159, the flanking regions (upstream fragment of 635 bp, downstream fragment of 574 bp) were fused with an erythromycin cassette into a single PCR product. For transformation, *S. mutans* was grown in BHI supplemented with heat-inactivated horse serum and the PCR fusion construct was added at 0.5 μg/mL.

Complementation of both *S. aureus* (MRSA Δ*lspA*::p*lspA*) and *S. mutans* strains was performed with pDC123 containing the full length *lspA* (SAUSA300_1089 for *S. aureus* or SMU_853 for *S. mutans*, respectively). Successful transformation was checked with chloramphenicol resistance and colony PCR. An overview of all strains, plasmids and primers used in this study are shown in Tables 1 and 2. All transformants were plated on selective plates containing appropriate antibiotics and successful transformation was checked with PCR and sequencing.

**Table 2.**
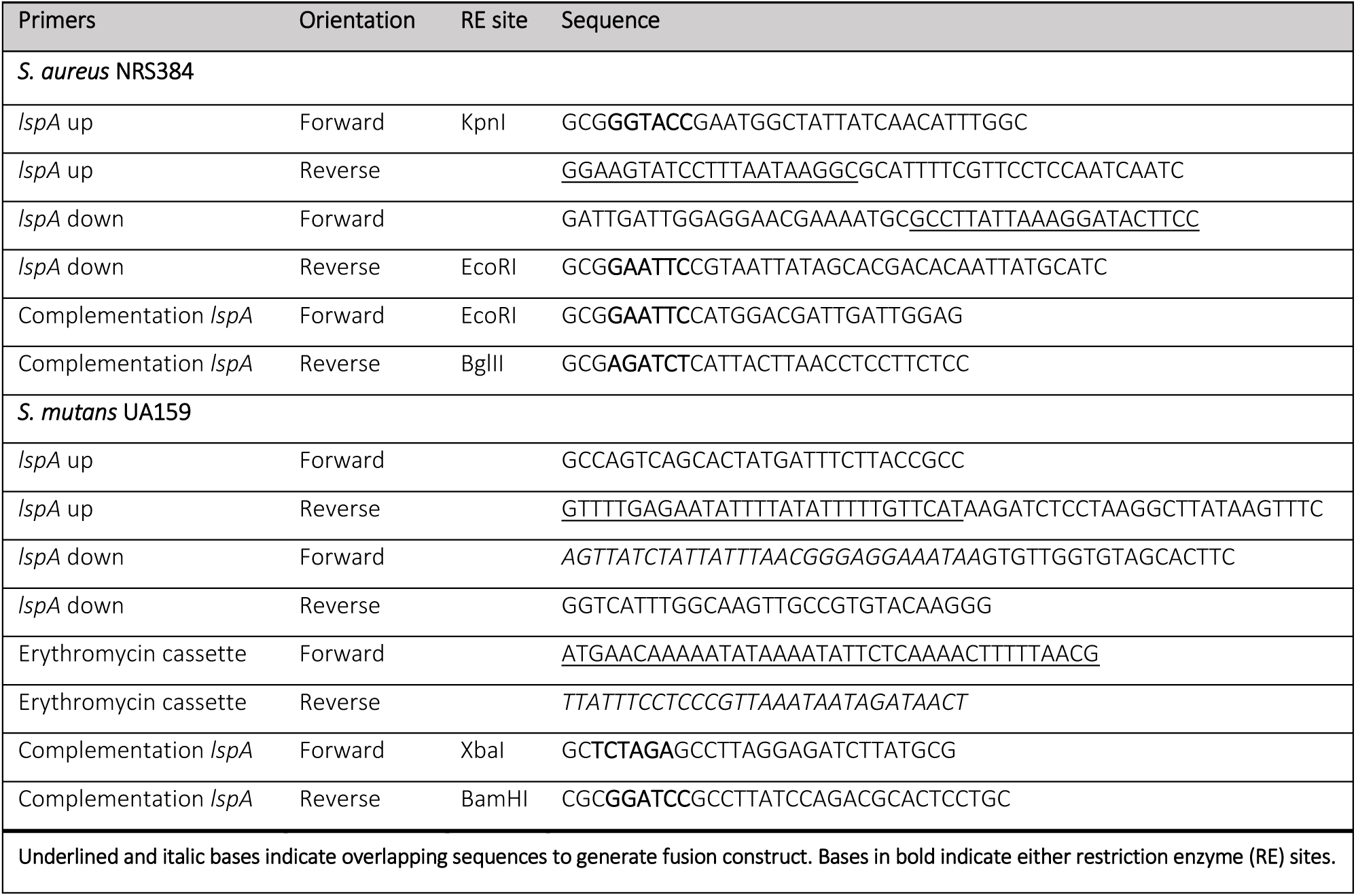
Overview of primers used in this study.

### CFU killing assay

Survival after hGIIA, daptomycin, or gentamicin exposure was determined by quantifying CFU on TH agar. Early log-phase bacteria (OD_600_ of 0.2 for S. *mutans* or 0.4 for *S. aureus* and *Enterococcus* spp.) were washed and resuspended in HEPES solution supplemented with 1% BSA (HEPES 1% BSA) and cell density was adjusted to the original OD_600_. Bacterial suspensions (containing 10^3^ CFU of *S. aureus*, 2×10^3^ CFU of *S. mutans* or 10^5^ CFU of *Enterococcus* spp.) were mixed 1:1 with increasing concentrations of recombinant hGIIA, daptomycin, or gentamicin in HEPES 1% BSA and incubated for 1 hour at 37°C. Samples were then serially diluted in phosphate buffered saline (PBS, pH 7) and plated on TH agar plates. After overnight incubation at 37°C, CFU were counted and bacterial survival was calculated compared to untreated bacteria. To investigate the effect of the LspA inhibitor globomycin or myxovirescin A1 on hGIIA- or daptomycin-mediated killing, the compounds were added to wild-type (WT) bacteria during sub-culturing to early exponential phase at a concentration of 100 μg/mL for globomycin and 10 μg/mL for myxovirescin A1, which were produced and purified as previously described [44] and dissolved in DMSO. The maximum concentration of DMSO was 1%, which was also added to other bacterial cultures as a control.

### Scanning Electron Microscopy (SEM)

MRSA WT, MRSA Δ*lspA*, and MRSA Δ*lspA*::p*lspA* at stationary phase and early exponential phase (OD_600_ 0.4) were washed, fixed, and dehydrated as described previously [45]. Samples were mounted on 12.5 mm specimen stubs (Agar scientific, Stansted, Essex, UK) and coated with 1 nm gold using the Quorum Q150R S sputter coater at 20 mA. Microscopy was performed with a Phenom PRO desktop SEM (Phenom-World BV) operating at an acceleration voltage of 10 kV.

### Growth curve

MRSA WT, MRSA Δ*lspA*, and MRSA Δ*lspA*::p*lspA* were grown overnight and sub-cultured the following day to an OD_600_ of 0.4 in THB supplemented with antibiotics when appropriate. The early exponential phase bacteria were diluted to OD_600_ 0.025 in THB. OD_600_ was measured every 5 minutes over 20 hours (shaking) in a Biotek Synergy H1.

### MRSA infection experiment in hGIIA-Tg mice

Tg mice overexpressing hGIIA were from Taconic (Denmark). They were generated by inserting the 6.2 kb full-length of human gene (*PLA*_*2*_*G2A*) into the mouse genome and were bred to a sPLA_2_-IIA naturally-deficient C57BL/6 female mouse that lacks the functional mouse homologue (Pla2g2a) [19, 46]. The animals were housed at Institut Pasteur animal facility accredited by the French Ministry of Agriculture for performing experiments on live rodents. The study on animals was performed in compliance with the French and European regulations on care and protection of laboratory animals (EU Directive 2010/63, French Law 2013-118, February 6th, 2013). The experimental protocol was approved by the Institut Pasteur Ethics Committee and registered under the reference 2014–0014 with the infection protocol 21.185 (AC 0419).

Mice, both males and females (Supplementary Table 1) of 7–9 weeks old, were bred at Institut Pasteur animal facility and infected intra-peritoneally with MRSA WT or the isogenic Δ*lspA* mutant (1×10^7^ or 5×10^7^ CFU) suspended in 100 μL PBS. Mortality and weight loss of mice were monitored twice daily up to 5 days after infection.

### Surface charge

Bacterial surface charge was determined as previously described [47]. Briefly, early-exponential phase bacteria (OD_600_ = 0.4) were washed twice in 20 mM MOPS buffer (pH 7.0, Sigma-Aldrich) and adjusted to OD_600_ 0.7. Bacteria were concentrated 10 times, of which 200 μL aliquots were added to 0.5 mg/mL cytochrome c (from Saccharomyces cerevisiae, Sigma-Aldrich) in a sterile 96 well round-bottom plate. Suspensions were incubated for 10 minutes at room temperature and subsequently centrifuged at 3,500 rpm for 8 minutes. Supernatant was transferred to a sterile 96 well flat-bottom plate and absorbance was recorded at 530 nm. The percentage of residual cytochrome c was calculated using samples containing MOPS buffer only (100% binding) and samples containing MOPS buffer and cytochrome c (0% binding).

### Membrane potential and permeability assays

Changes in hGIIA-dependent membrane potential were determined using the membrane potential probe DiOC_2_(3) (PromoKine) [15, 48]. Bacterial suspensions (OD_600_ of 0.4) were diluted 100 times (∼1×10^6^ CFU/mL) and incubated with serial dilutions of hGIIA. After incubation at 37°C, 3 mM DiOC_2_(3) was added and incubated at room temperature for 5 minutes in the dark. Changes in green and red fluorescence emissions were analyzed by flow cytometry. Bacterial staining with the DNA stain SYTOX Green (Invitrogen) is a measurement for membrane permeabilization and an indication of bacterial cell death [49]. Serial dilutions of hGIIA in HEPES solutions were added to wells of a sterile flat-bottom 96 well plate. Bacteria were resuspended in HEPES solution containing 1 μM SYTOX green (OD_600_ of 0.4) and added to hGIIA dilutions in a final volume of 100 μL. Fluorescence over time was recorded using Optima Fluostar (green fluorescence 520 nm emission and excitation 485 nm) at 37°C.

### PubMLST database analysis of *S. aureus lspA*

The PubMLST database, assessed at https://pubmlst.org/organisms/staphylococcus-aureus [50] was used to analyze the presence and sequence conservation of *lspA* (SAUR1197) across the *S. aureus* population. Alignments were made using the locus explorer of the PubMLST database and nucleotide and amino acid identity was calculated using the NCBI BLAST tool (https://blast.ncbi.nlm.nih.gov/blast.cgi). *LspA* gene sequences of 26,036 *S. aureus* strains were downloaded from the database in February 2021. We excluded whole genome sequences for data analysis that were unlikely to be *S. aureus*, contained > 300 contigs or an N50 contig length shorter than 20,000 bp, contained an internal stop codon rendering a truncated LspA or when *lspA* was located at the end of a contig.

### Statistical analysis

Statistical analysis was performed using GraphPad Prism 9. We used the Student’s *t* test and one- and two-way ANOVA’s with Bonferroni statistical hypothesis testing to correct for multiple comparisons. All values are reported as mean with standard error of the mean of three biological replicates unless indicated otherwise. A *p* value of < 0.05 was considered statistically significant.

## Results

### Identification of hGIIA resistance genes in MRSA

To unravel new hGIIA resistance mechanisms of MRSA, we screened 1,920 individual MRSA mutants of the Nebraska Transposon Mutant Library (NTML). Exponentially-grown transposon mutants were exposed to recombinant hGIIA for one hour and subsequently spotted on agar plates for semi-quantitative assessment of survival (Supplementary Figure 1A). In total, 39 mutants were identified with potential increased susceptibility to hGIIA-mediated killing (Supplementary Table 2). These hits included the transposon mutant NE1360 (*mrpF*), which displays an increased positive charge of membrane phospholipids and was previously linked to hGIIA resistance [14]. Additionally, transposon insertion in genes encoding the two-component system GraRS and its ABC-transporter VraFG also rendered MRSA more susceptible to hGIIA. These genes are important for the regulation of the before mentioned *mprF* and *dltABCD* operon [22], which has a known role in hGIIA resistance [14]. Transposon mutants in individual genes of the *dltABCD* operon were not identified since these mutants are absent in the NTML [34].

To confirm the phenotype of individual transposon mutants identified in our screen, we assessed their susceptibility in a quantitative killing assay across a hGIIA concentration range. As expected, disruption of previously-identified genes *graR, graS*, and *mprF* rendered MRSA more susceptible to hGIIA-mediated killing (Supplementary Figure 1B). In contrast, mutants with transposons inserted in the genes *esaC, srtB, ltaA*, and *asp1* were not differently affected by hGIIA (Supplementary Figure 1B). Interestingly, the *lspA* transposon mutant (NE1757), showed increased susceptibility to hGIIA (Supplementary Figure 1B). *LspA* is conserved among Gram-positive and Gram-negative bacteria and encodes the lipoprotein signal peptidase A, an enzyme involved in the lipoprotein maturation pathway [30, 51].

### Deletion of *lspA* attenuates MRSA resistance to hGIIA *in vitro* and virulence in a hGIIA-Tg mouse model

To verify the contribution of LspA to hGIIA resistance, we constructed a markerless *lspA* deletion mutant in the MRSA strain NRS384 (MRSA Δ*lspA*) and a plasmid complemented mutant strain (MRSA Δ*lspA*::p*lspA*). In accordance to results from our NTML screen, MRSA Δ*lspA* was 5 to 10-fold more susceptible to hGIIA-mediated killing and the phenotype was rescued by complementation with the full length *lspA* gene (Fig. 1A). Deletion of *lspA* in the MRSA background did not result in morphological differences as assessed by scanning electron microscopy (Fig. 1B). Moreover, in accordance with previous literature of other Gram-positive bacteria [52-55], growth of MRSA in bacterial broth was not affected in the *lspA* deletion mutant (Fig. 1C).

**Figure 1.**
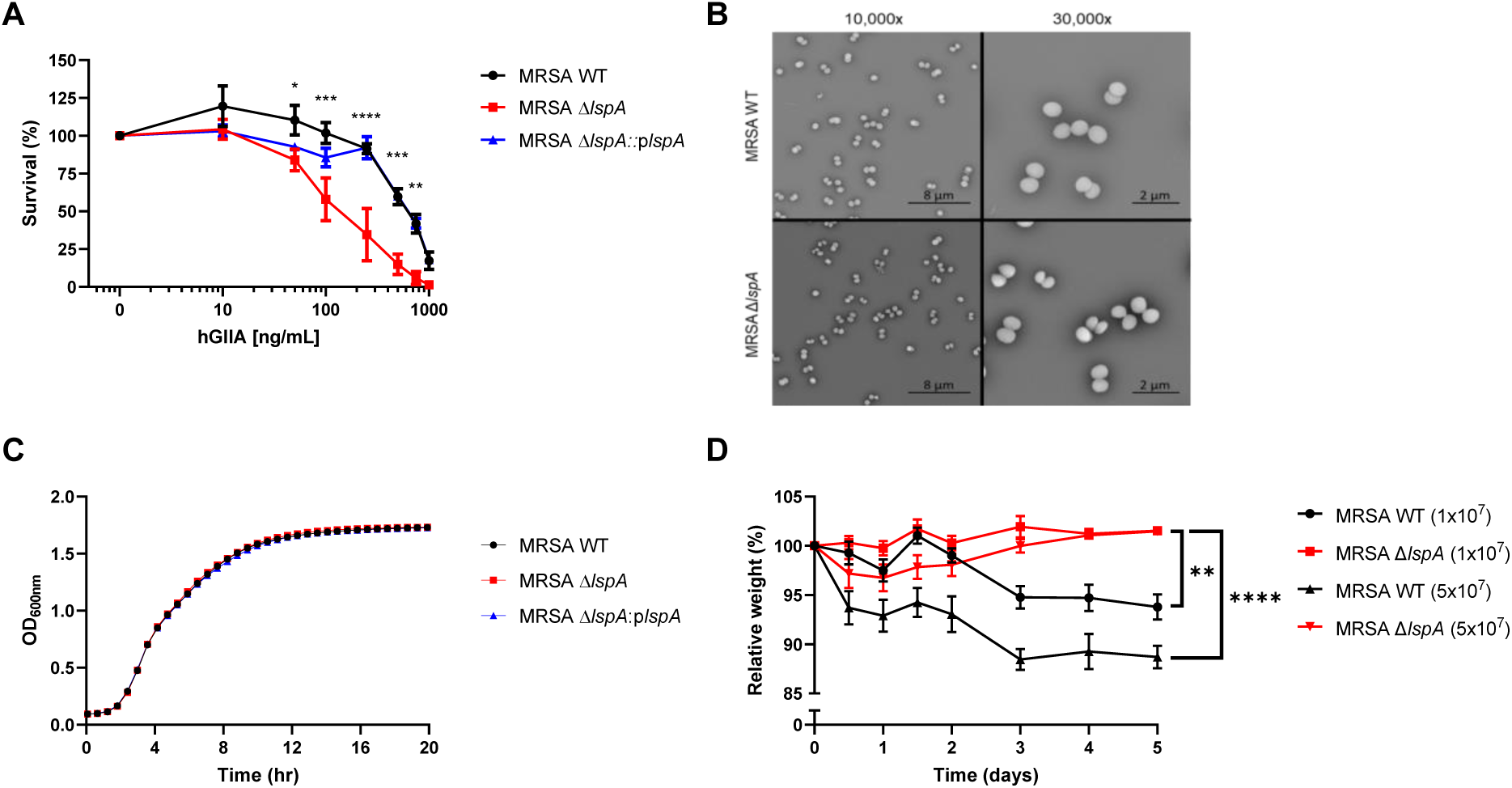
LspA contributes to hGIIA resistance *in vitro* as well as virulence in a hGIIA-Tg mouse model. (A) Survival of MRSA WT, MRSA Δ*lspA*, and MRSA Δ*lspA*::p*lspA* after exposure to a concentration range of recombinant hGIIA. (B) Representative scanning electron microscopy (SEM) images of MRSA WT and MRSA Δ*lspA* in early exponential phase. (C) Growth curves of MRSA WT, MRSA Δ*lspA*, and MRSA Δ*lspA*::p*lspA*. (D) Relative weight of male and female hGIIA-Tg C57BL/6 mice injected i.p. with either MRSA WT or MRSA Δ*lspA* (1×10^7^ or 5×10^7^ CFU). Statistical significance was determined using one- or two-way ANOVA + Bonferroni’s Multiple Comparison Test. \**p* < 0.05, \*\**p* < 0.01, \*\*\**p* < 0.001, \*\*\*\**p* < 0.0001. A, D: Data represent mean with standard error of the mean of three biological replicates.

It was previously shown that mutation of *lspA* resulted in attenuated virulence of *S. aureus* but had no effect on median lethal dose (LD_50_) values in a mouse infection model [31]. Interestingly, the mouse strain used in this study was C57BL/6, which lacks a functional mouse sPLA_2_-IIA homologue due to a natural frameshift mutation [19]. Although hGIIA-Tg mice, generated in this naturally-deficient strain background, showed enhanced survival compared to control littermates after infection with WT *S. aureus* [18], it has not yet been determined how *lspA* mutation affects *S. aureus* virulence in a mouse strain with a functional GIIA gene. Therefore, we infected hGIIA-Tg C57BL/6 mice with MRSA WT or its isogenic mutant Δ*lspA* at 2 different doses (1×10^7^ or 5×10^7^ CFU/mouse). All mice survived the challenge. However, as judged by weight loss, mice infected with either 1×10^7^ or 5×10^7^ MRSA WT bacteria showed significantly more weight loss compared to mice infected with Δ*lspA* bacteria (Fig. 1D). This suggests that in the presence of a functional hGIIA enzyme, LspA contributes to MRSA virulence in this infection model.

### hGIIA shows faster cell wall penetration and membrane permeabilization in the absence of LspA

To gain further insights into the underlying mechanisms of hGIIA susceptibility in the absence of LspA, we assessed the effects of *lspA* deletion on hGIIA binding and cell wall penetration. Since charge-dependent binding is an important first step in hGIIA’s mechanism of action, we determined the surface charge of the three strains using the cationic compound cytochrome c [47]. Equal binding levels of cytochrome c was observed for all three strains (Fig. 2A), suggesting that *lspA* does not affect surface charge. However, we did observe that MRSA Δ*lspA* was not only more sensitive to hGIIA, but that killing kinetics were also faster for the mutant compared to WT (Fig. 2B). To assess whether hGIIA trafficking across the cell wall was different, we compared how hGIIA affected membrane depolarization (early effect of hGIIA activity) and membrane permeabilization (late effect of hGIIA activity). Membrane depolarization was measured with the fluorescent voltage-sensitive dye DiOC_2_(3) that exhibits green fluorescence (FITC) in all bacterial cells dependent on cell size and red fluorescence (PerCP) dependent on membrane potential. Deletion of *lspA* resulted in a faster and more extensive membrane depolarization (Fig. 2C, Supplementary Figure 2). Loss of LspA also caused increased SYTOX intensity, an indication of membrane permeabilization [15, 47], compared to MRSA WT and complemented strain starting from 9 min (Fig. 2D).

**Figure 2.**
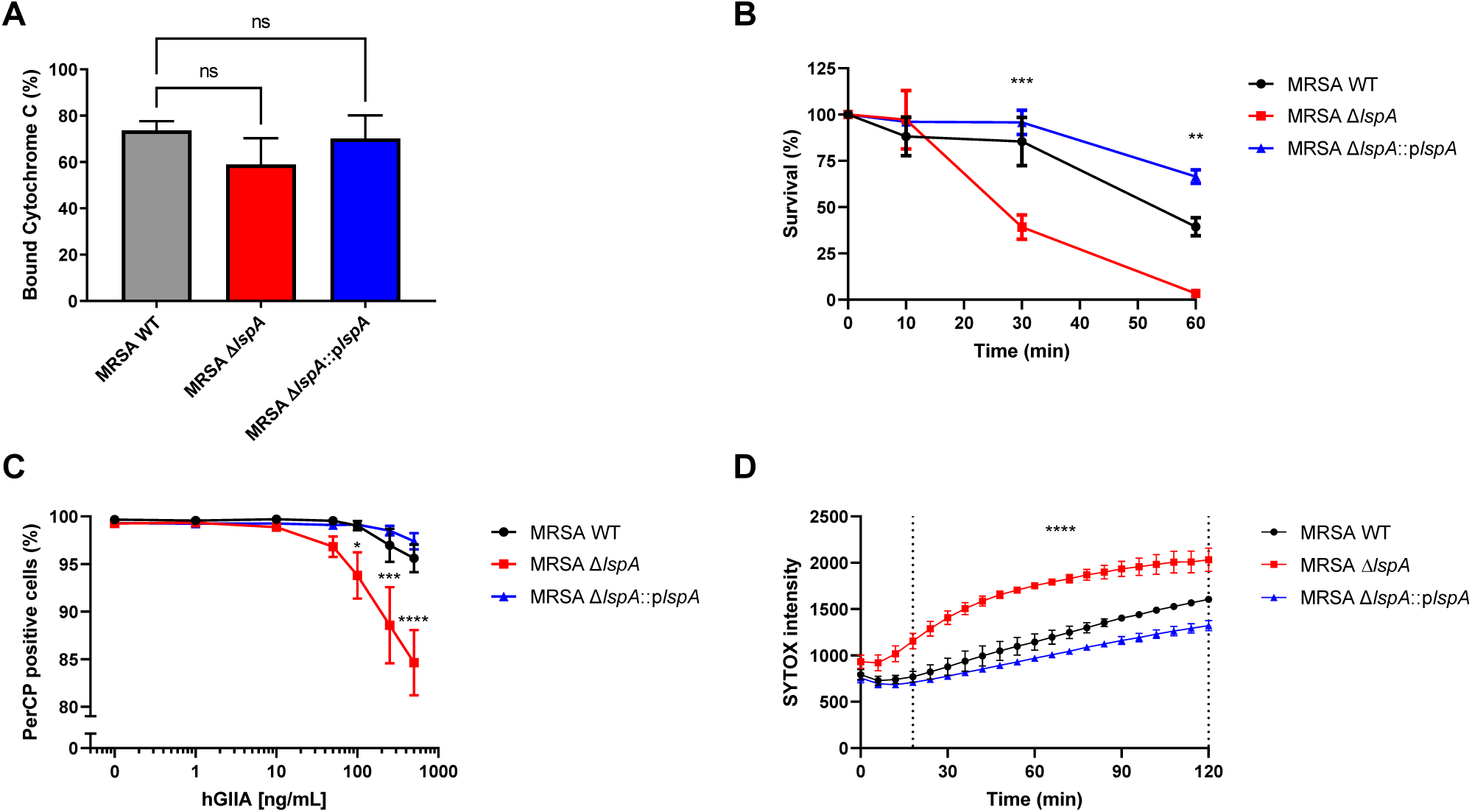
Deletion of *lspA* results in faster hGIIA cell wall penetration and membrane permeabilization. (A) Surface charge of MRSA WT, MRSA Δ*lspA*, and MRSA Δ*lspA*::p*lspA* as determined in a cytochrome c binding assay. (B) Survival of MRSA WT, MRSA Δ*lspA*, and MRSA Δ*lspA*::p*lspA* over time after incubation with 500 ng/mL recombinant hGIIA. (C) Flow cytometric analysis of PerCP-positive cells of MRSA WT, MRSA Δ*lspA*, and MRSA Δ*lspA*::p*lspA* stained with DiOC_2_(3) after exposure to a concentration range of recombinant hGIIA. (D) Kinetic analysis of SYTOX intensity for MRSA WT, MRSA Δ*lspA*, and MRSA Δ*lspA*::p*lspA* in the presence of 250 ng/mL recombinant hGIIA. Statistical significance was determined using a one- or two-way ANOVA + Bonferroni’s Multiple Comparison Test. ns = not significant, \**p* < 0.05, \*\**p* < 0.01, \*\*\**p* < 0.001, \*\*\*\**p* < 0.0001. Data represent mean with standard error of the mean of three biological replicates.

### Interruption of lipoprotein maturation sensitizes MRSA towards daptomycin

The antibiotic daptomycin is clinically important to treat MRSA infections. Interestingly, the mechanism of action of daptomycin displays similarities with hGIIA, since it is dependent on its positive charge and targets the cell membrane [9, 56]. Correspondingly, the identified *S. aureus* resistance genes, i.e. *dltABCD, graRS*, and *mprF* overlap for daptomycin and hGIIA [14, 22-24, 57]. We therefore investigated whether *lspA* deletion affected daptomycin resistance. Indeed, MRSA Δ*lspA* was about 5-fold more susceptible to daptomycin killing, whereas the *lspA* plasmid complemented strain became even more resistant compared to WT (Fig. 3A). As comparison, we assessed whether an intracellular acting antibiotic, gentamicin, was differentially effective in the presence and absence of LspA. Only at one concentration did we observe that loss of *lspA* rendered MRSA more susceptible to gentamicin killing (Fig. 3B), indicating that LpsA has minimal impact on gentamicin-mediated killing.

**Figure 3.**
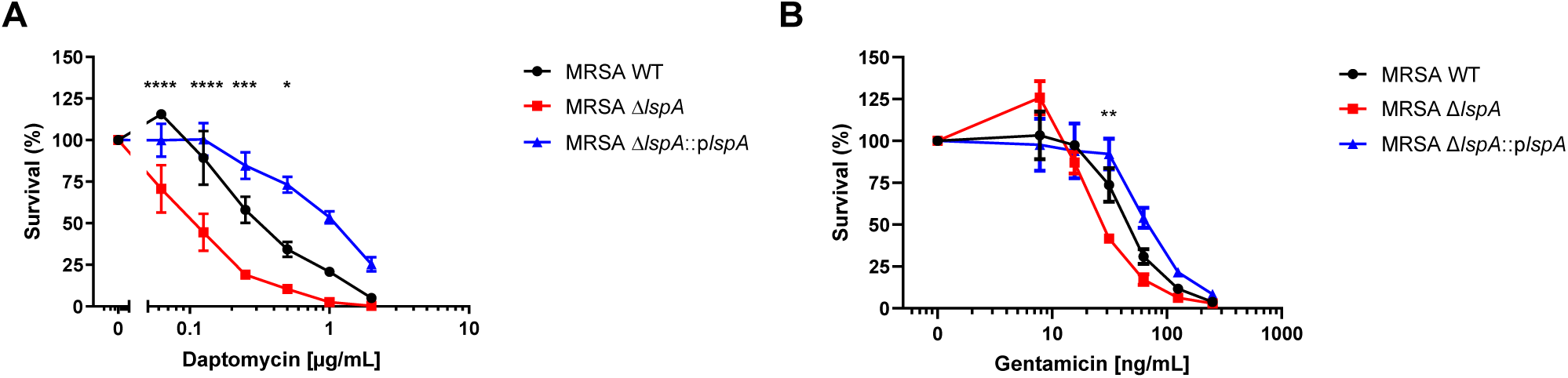
Impact of LspA on killing by clinically-relevant antibiotics. Survival of MRSA WT, MRSA Δ*lspA*, and MRSA Δ*lspA*::p*lspA* after exposure to (A) daptomycin or (B) gentamicin. Statistical significance was determined between MRSA WT and MRSA Δ*lspA* using a two-way ANOVA + Bonferroni’s Multiple Comparison Test. \**p* < 0.05, \*\**p* < 0.01, \*\*\**p* < 0.001, \*\*\*\**p* < 0.0001. Data represent mean with standard error of the mean of three biological replicates.

### LspA inhibitors sensitize MRSA towards hGIIA and daptomycin

The antibiotics globomycin and myxovirescin A1 are directly bactericidal towards Gram-negative bacteria with minimum inhibitory concentration values of 12.5 and 1 μg/mL for *E. coli*, respectively [58, 59]. Interestingly, both compounds are LspA inhibitors [60, 61] and do not kill *S. aureus* growth even at concentrations of 30 μg/mL myxovirescin A1 and >100 μg/mL globomycin [58, 59]. The co-crystal structures of *S. aureus* LspA with both of these inhibitors were recently published [44]. We assessed whether MRSA could also be sensitized to hGIIA and daptomycin through pharmacological inhibition of LspA. To this end, we pre-incubated MRSA WT with either of these compounds during growth to exponential phase and subsequently exposed the bacterial culture to hGIIA or daptomycin. Indeed, pharmacological interference with LspA by either compound rendered MRSA more susceptible to killing by hGIIA and daptomycin compared to untreated bacteria (Fig. 4A, B). This suggests that these compounds may be interesting sensitizing agent in the context of *S. aureus* infections.

**Figure 4.**
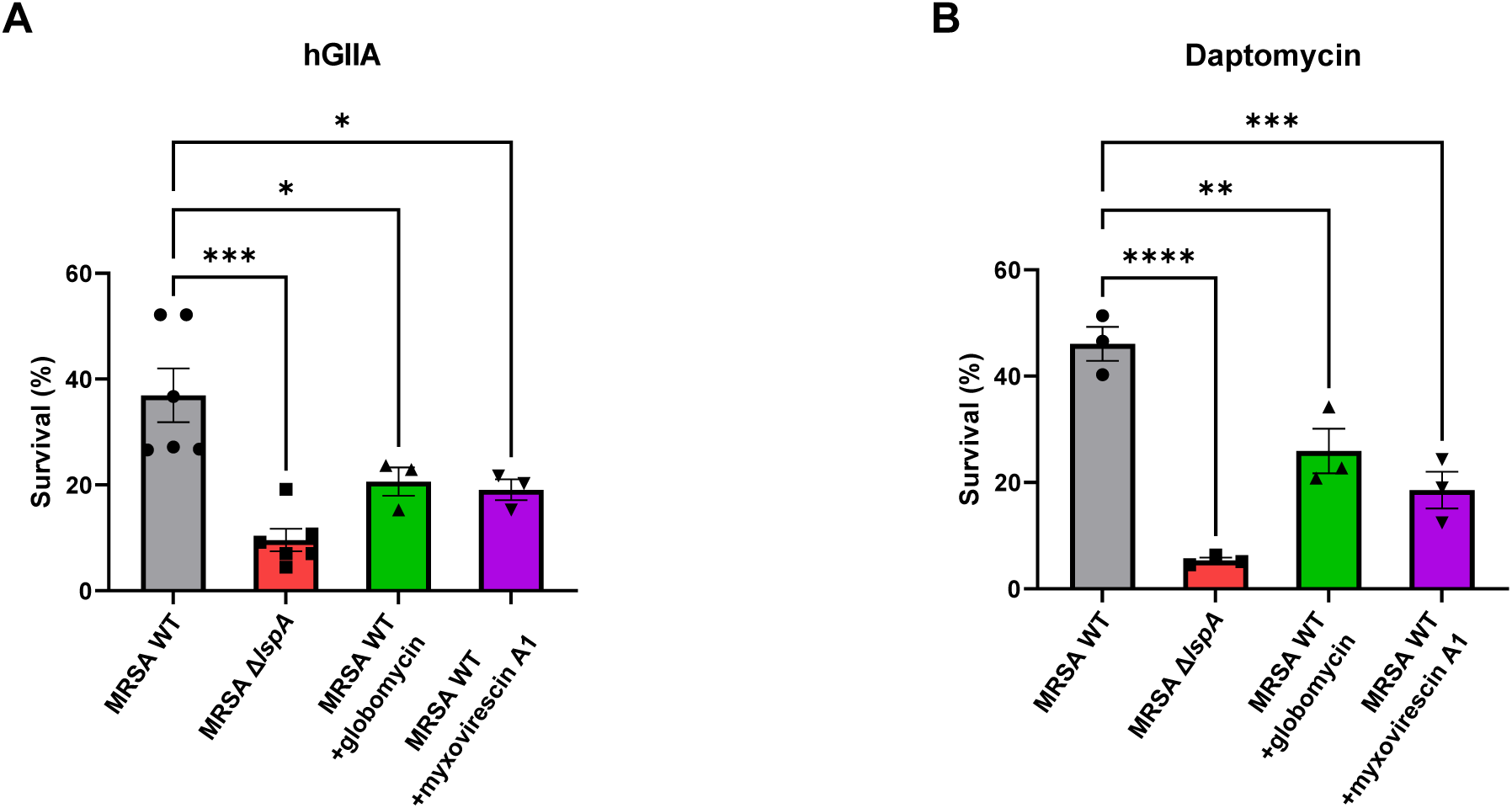
Globomycin and myxovirescin A1 increase MRSA killing by hGIIA and daptomycin. Survival of MRSA WT, MRSA Δ*lspA*, MRSA WT + 100 μg/mL globomycin, and MRSA WT + 10 μg/mL myxovirescin A1 after subsequent exposure to (A) recombinant hGIIA (250 ng/mL) or (B) daptomycin (1 μg/mL). Statistical significance was determined using a one-way ANOVA + Bonferroni’s Multiple Comparison Test. \**p* < 0.05, \*\**p* < 0.01, \*\*\**p* < 0.001, \*\*\*\**p* < 0.0001. Data represent mean with standard error of the mean of three biological replicates.

### LspA is highly sequence-conserved within the *S. aureus* population

In considering LspA as a drug target, it is important to assess the sequence conservation over bacterial species. LspA contains five conserved domains, including the catalytic residues, across several bacteria [62]. Moreover, LspA amino acid sequence identity is in-between 35% and 95% across 485 different bacterial species [63].

To investigate the presence and sequence conservation of *lspA*, the genomes of 25,243 *S. aureus* isolates were surveyed using PubMLST [50]. These isolates originated from different continents and from a wide variety of hosts as well as human patients and carriers. A *lspA* gene was present in all isolates examined. Only 5 isolates contained a gene with an internal stop codon rendering a truncated LspA. In total, 141 *lspA* alleles were observed.

The majority of the isolates contained *lspA*4 (14,000 isolates, 54%), *lspA*5 (6,000 isolates, 23%) or *lspA*1 (4,000 isolates, 16%). All other *lspA* alleles were at frequencies < 2.5% (Table 3). Interestingly, specific clonal complexes were associated with a single dominant allele (Table 3). Among 141 *lspA* alleles 110 polymorphic positions out of a total gene length of 492 nucleotides were found. These 110 polymorphic sites represented 124 single nucleotide polymorphisms (SNPs). None of the SNPs found in critical residues were synonymous, emphasizing the high degree of conservation. The most frequently observed SNPs were found at nucleotide positions 331 and 402 (Fig. 5A). Only one of these, at nucleotide position 331, and present in *lspA*1, *lspA*3, *lspA*7 and *lspA*26, results in an amino acid substitution (Ile111Val). A second non-synonymous SNP at nucleotide position 230 is found in *lspA*3, but this allele is present in only 1% of the isolates (Table 3 and Fig. 5B). All other SNPs, as found in the most frequently observed *lspA* alleles among the *S. aureus* population studied are synonymous (Fig. 5B).

**Table 3.** Distribution of *lspA* among 25,243 S. *aureus* isolates.

| *lspA* allele | # isolates | Percentage <sup>a</sup> | Dominant in cc <sup>b</sup> |
| --- | --- | --- | --- |
| 4 | 14,000 | 53.7% | 1, 8, 15, 22, 97 |
| 5 | 6,000 | 22.6% | 5 |
| 1 | 4,000 | 15.7% | 30 |
| 7 | 500 | 2.2% | 45 |
| 26 | 400 | 1.7% | 93 |
| 3 | 300 | 1.1% | - |
| Other | 700 | 2.9% |  |
| <sup>a</sup>Alleles with at least 1% occurrence in isolates are shown<br><sup>b</sup>The allele was considered most dominant in the clonal complex (cc) with an occurrence percentage of 92% or higher.<br>Numbers are rounded off to thousands and to tenths for number of isolates and percentage, respectively. |  |  |  |

**Figure 5.**
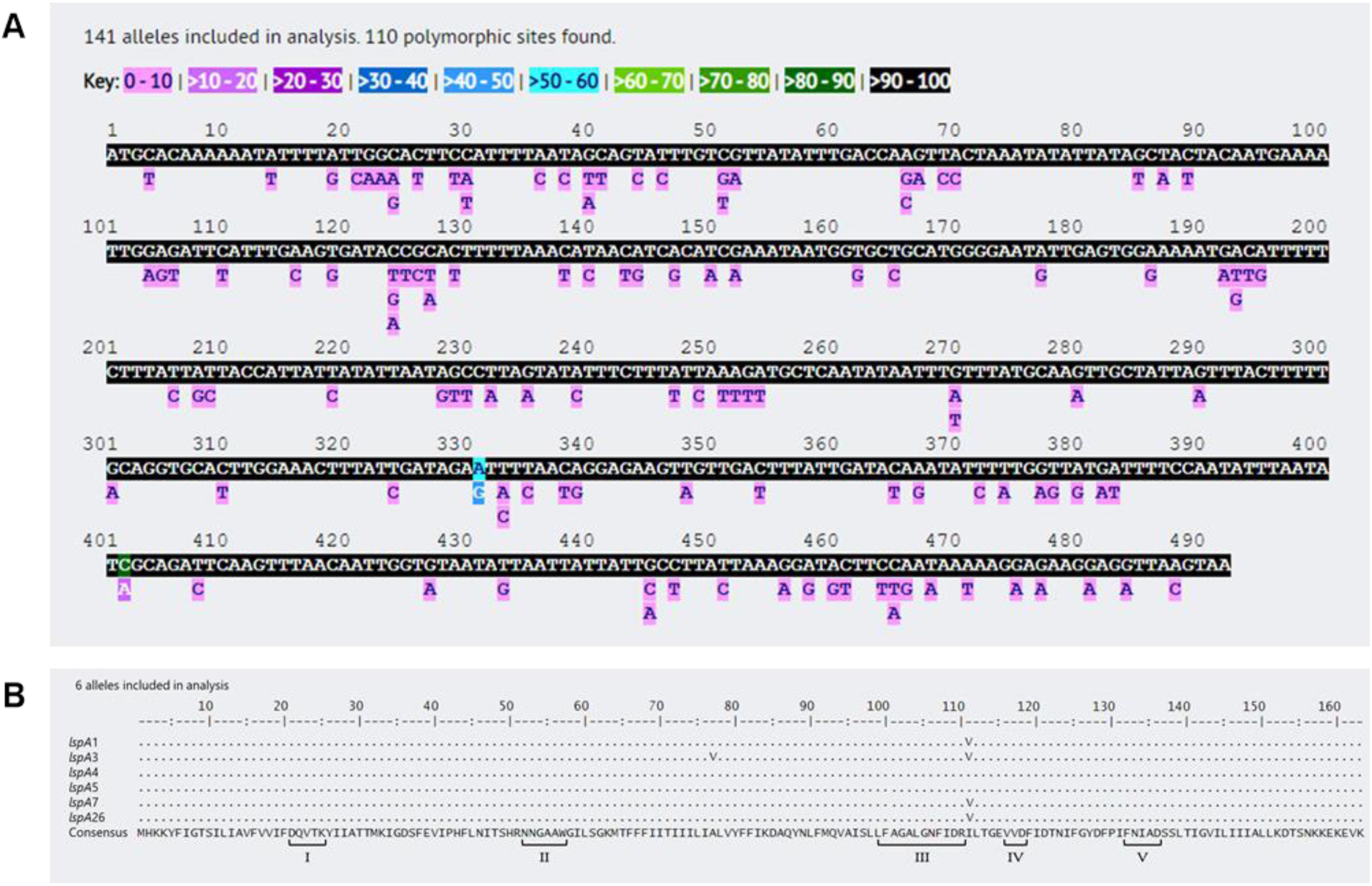
*lspA* and the encoded LspA protein are highly sequence conserved across the *S. aureus* population. (A) Polymorphic site frequencies of 141 alleles of *lspA* among 25,243 *S. aureus* genomes. Consensus sequence is depicted with color coding for the occurrence in percentages. (B) Alignment and consensus sequence at amino acids level encoded by the 6 most common *lspA* alleles. The five conserved domains across all bacterial species are depicted below with roman numerals [62]. *LspA*4 is the reference allele.

Thus, only two amino acid differences are found when comparing the protein sequences encoded by the six most frequently found alleles among a total population of 25,243 isolates analyzed.

### LspA contribution to hGIIA resistance is not restricted to *S. aureus*

*Streptococcus mutans* is a Gram-positive bacterium that resides in the human oral cavity and is the major cause of dental caries [64]. To assess whether LspA-mediated resistance to hGIIA is restricted to

*S. aureus* or more widespread, we created a *lspA* deletion mutant in *S. mutans* strain UA159 by replacing the *lspA* gene with an erythromycin cassette. Complementation of this deletion mutant was accomplished by introducing the plasmid pDC123 containing the full *lspA* gene of *S. mutans*. Results from the killing assay revealed that *lspA* deletion renders *S. mutans* more susceptible to hGIIA and complementation fully restored this phenotype (Fig. 6A).

**Figure 6.**
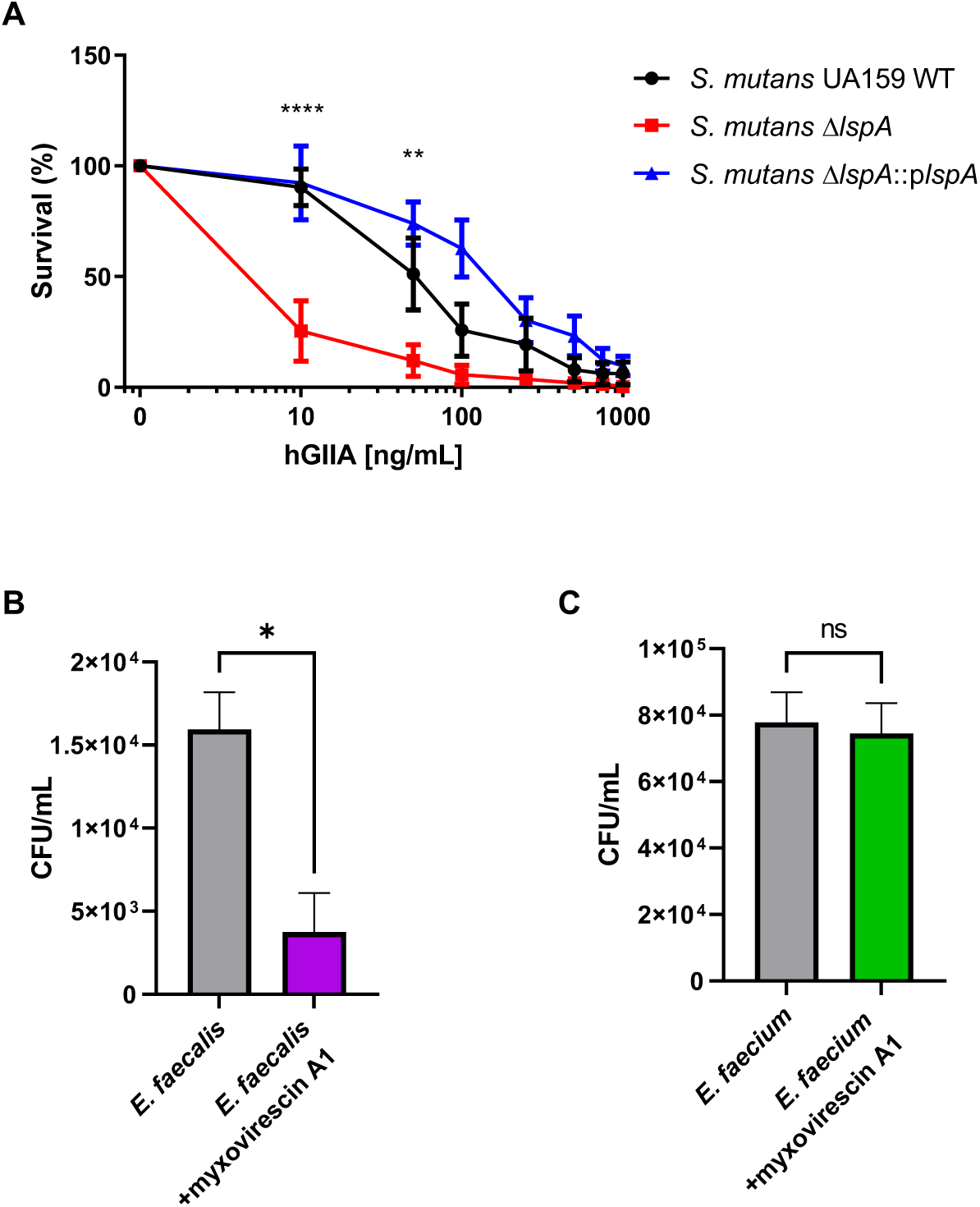
*S. mutans* and *E. faecalis*, but not *E. faecium*, are sensitized to hGIIA via *lspA* deletion or LspA inhibition. (A) Survival of *S. mutans* UA159 WT, *S. mutans* Δ*lspA*, and *S. mutans* Δ*lspA*::p*lspA* after exposure to concentration range of recombinant hGIIA. (B) Survival of *E. faecalis* and *E. faecalis* + 10 μg/mL myxovirescin A1 after subsequent exposure to recombinant hGIIA (0.5 ng/mL). (C) Survival of *E. faecium* and *E. faecium* + 10 μg/mL myxovirescin A1 after subsequent exposure to recombinant hGIIA (0.5 ng/mL). Statistical significance was determined using a one- or two-way ANOVA + Bonferroni’s Multiple Comparison Test or an unpaired two-tailed Student’s *t* test. ns = not significant, \**p* < 0.05, \*\**p* < 0.01, \*\*\*\**p* < 0.0001. Data represent mean with standard error of the mean of three biological replicates.

In addition, we tested two clinical isolates of the enterococcal strains *E. faecalis* V583 and *E. faecium* U0317. These species are part of a group that consists of clinically-relevant and antibiotic-resistant pathogens, collectively called ESKAPE pathogens (*Enterococcus* spp., *Staphylococcus aureus, Klebsiella pneumoniae, Acinetobacter baumannii, Pseudomonas aeruginosa* and *Enterobacter* spp.) [65, 66]. Of the Gram-positive *Enterococci*, the species *E. faecalis* and *E. faecium* are most abundant and are responsible for 75% of all enterococcal infections [67]. We observed that *E. faecalis* was 5-fold more sensitive to hGIIA compared to *E. faecium* (Supplementary Figure 3). Pretreating the clinical enterococcal isolates with 10 μg/mL myxovirescin A1 sensitized *E. faecalis*, but not *E. faecium*, to hGIIA killing compared to the untreated bacteria (Fig. 6B, C). Also, higher concentrations of myxovirescin A1 (i.e. 50 μg/mL) did not increase hGIIA killing of *E. faecium*.

## Discussion

New treatment strategies against MRSA are in high demand due to the rise of antibiotic resistance even against the last-resort antibiotic daptomycin. The current antibiotic arsenal as well as many therapeutic agents under development aim to be directly bactericidal or stop bacterial growth [6]. The drawback of these compounds is the high selective pressure leading to antimicrobial resistance. Non-traditional antibacterial agents, such as anti-virulence drugs, can offer new therapies in the race against antimicrobial resistance by interfering with bacterial strategies that normally allow survival in the context of immune defenses [68]. Such strategies are expected to be less affected by resistance development as there is no direct pressure on survival [69]. Although sensitizing agents still have to prove their clinical use, the concept is appealing. Many of these strategies against *S. aureus* are under active investigation and some are already in preclinical development [70]. For example, inhibition of staphyloxanthin production increased susceptibility to killing in human blood and decreased the virulence of *S. aureus* in mouse infection models [71, 72]. The present work shows that interfering with lipoprotein maturation by inhibition of LspA enhances innate immune killing of MRSA through the modulation of the bactericidal effects of hGIIA. LspA inhibition also enhances daptomycin-mediated killing, which may provide an add-on strategy in antibiotic treatment.

To identify resistance genes against hGIIA in MRSA, we screened the NTML and confirmed increased susceptibility for hits in *graR, graS*, and *mprF*. These three genes have previously been linked to cationic antimicrobial resistance [22], and also specifically to hGIIA resistance [14]. We also identified *vraF* and *vraG*, which is also in line with expectations, since these genes encode the ABC-transporter linked to the GraRS two-component system [73]. This confirms that the screen, although semi-quantitative, does allow the identification of hGIIA-susceptible mutants. However, the screen likely lacks sensitivity to provide a comprehensive list of hGIIA-susceptible mutants. This is illustrated by the fact that we did not identify *graX*, the gene encoding GraX, which was shown to be involved in cationic antimicrobial peptide resistance and interacts with the GraRS system [73, 74]. Therefore, additional hGIIA sensitive mutants are likely to be identified using another set-up of the screening assay.

In our unbiased genetic screen, we identified the transposon mutant NE1757 (*lspA*) to be more susceptible to hGIIA-mediated killing. To exclude the possibility that the *lspA* transposon mutant was identified as a result of growth defects or polar effects of the transposon insertion, we constructed a *lspA* deletion strain in the MRSA background NRS384 that was exposed to a hGIIA concentration range and quantified for bacterial survival. With this quantitative killing assay as well as an infection model in hGIIA-Tg mice, we confirmed *lspA* as a novel hGIIA resistance determinant. Additionally, MRSA Δ*lspA* was also more effectively killed by daptomycin compared to WT. This makes LspA an interesting therapeutic target as its inhibition would simultaneously increase susceptibility to endogenous and specific clinically-used antibiotics.

Indeed, we provided proof-of-principle that inhibition of LspA by two known pharmacological inhibitors, globomycin and myxovirescin, renders MRSA more susceptible to hGIIA and daptomycin killing. A previous study has shown that the *S. aureus* LspA enzyme is inhibited by these compounds, but has no direct bactericidal effects [44]. This is in line with the observation that deletion of *lspA* does not affect growth and morphological appearance of MRSA. Hence, selective pressure of this anti-virulence strategy is likely to be minimal. LspA inhibition as a therapeutic strategy may have other advantages. For example, the extracellular location of LspA makes it accessible to drug while no LspA analogs are found in eukaryotic cells, thereby reducing the risk of off-target effects [44, 62, 63]. In addition, we showed that LspA is highly conserved among *S. aureus* strains with only 1 amino acid substitution in >96% of the *S. aureus* collection in the PubMLST database (>26,000 isolates at the time of this analysis). Conserved proteins are less likely to mutate, making them ideal targets as the inhibitor compounds are longer lasting and more effective [75]. The natural antibiotics globomycin and myxovirescin A1 specifically inhibit LspA and have similar binding sites on LspA, docking to the catalytic dyad and clustering around 14 conserved residues [44, 60, 61, 63]. Although they have a distinct chemical structure and biosynthesis, there is a remarkable similarity in their mode of action. This might point towards a co-evolution that advanced to prevent resistance [44].

LspA processes prolipoproteins that are anchored into the cell membrane by the enzyme Lgt [30]. The mechanism by which LspA mediates hGIIA and daptomycin resistance is currently not clear. We explored the possibility that LspA deletion altered surface charge, thereby facilitating hGIIA binding. However, no difference in binding of the cationic protein cytochrome c was observed, suggesting no large effects on the net charge. Since hGIIA binding to bacteria is based on electrostatic interactions [76], we expect that hGIIA binds similar to WT and *lspA* knock-out strains. On the other hand, we did observe that loss of LspA affected both kinetics and concentration-dependent effects on membrane depolarization and membrane permeabilization, with Δ*lspA* mutants showing faster disruption after exposure to hGIIA. Since LspA is a transmembrane protein [62], lack of LspA may change membrane properties such as membrane fluidity. However, since we observed the same effects in MRSA WT after pretreatment with globomycin or myxovirescin, which inhibit LspA enzymatic activity, this explanation is unlikely. Nonetheless, the presence of multiple immature lipoproteins that still carry the signal peptide may affect membrane characteristics as these prolipoproteins likely accumulate in the membrane. In some Gram-positive bacteria other putative signal peptidases are present that could take over the role of LspA [26], but it is not known if this is the case in *S. aureus*. Another explanation could be that the function of a single lipoprotein is abolished by deletion of *lspA*, resulting in the observed phenotypes. However, our screen did not identify mutants in individual lipoprotein-encoding genes. In addition, lipoproteins may retain their function even without proper processing by LspA [77]. Based on these considerations and observations, we currently favor the hypothesis that differences in membrane composition due to the presence of the signal peptide are responsible for the observed phenotypes.

We observed that *lspA* deletion affected antibiotic susceptibility, most pronounced for daptomycin and marginally for gentamicin. In addition, daptomycin susceptibility could also be conferred by pharmacological inhibition of LspA. These findings suggest that LspA is involved in daptomycin resistance. However, the role of LspA in daptomycin-resistance is not necessarily straightforward, since *lspA* was not identified in two previous screens aimed at identifying daptomycin resistance determinants [78, 79]. The study using the same NTML as we did here [78], only identified a single daptomycin-susceptible mutant (SAUSA300_1003). This may indicate that the assay set up was unable to identify all susceptible mutants, since even *mprF*, a well-known daptomycin resistance determinant [24], was not identified. The second study used methicillin-sensitive *S. aureus* instead of MRSA to screen for antibiotic susceptibility, including daptomycin [79]. It may well be that strain background affects the contribution of *lspA* to daptomycin susceptibility. This is illustrated by a recent comparative transposon sequencing (Tn-seq) screen where only one of five *S. aureus* strains showed significant changes in *lspA* insertions after daptomycin exposure [80]. This observation may suggest that despite high protein sequence conservation, therapeutic efficacy of LspA inhibition may be strain-specific. This should be addressed in future studies when considering anti-virulence strategies.

Earlier *in vivo* experiments performed with a *S. aureus lspA* deletion strain showed that the mutant was less virulent [31, 32]. Interestingly, these experiments were performed in inbred C57BL/6 mice or outbred CD-1 mice, which carry a natural homozygous or heterozygous inactivating mutation in the mouse sPLA_2_-IIA-encoding gene, respectively [19]. Thus, to assess the contribution of *lspA* mutation to *S. aureus* virulence in an animal with a functional sPLA_2_-IIA enzyme, we performed a mouse infection experiment using hGIIA-Tg C57BL/6 mice [46]. These hGIIA-Tg mice have increased resistance to lethal *S. aureus* infection compared to control littermates [18]. In this hGIIA-Tg background, mice infected with MRSA Δ*lspA* did not display weight loss whereas mice infected with MRSA WT showed on average 5 to 10% weight loss depending on the infectious dose. Altogether, we conclude that LspA-dependent virulence occurs in a hGIIA-dependent and -independent manner as the effects are observed in naturally-deficient C57BL/6 mice and hGIIA-Tg mice.

The hGIIA susceptibility phenotype was not only observed in *S. aureus*, but also in *S. mutans* after *lspA* deletion or *E. faecalis* upon LspA inhibition. LspA inhibitors can bind LspA from multiple Gram-positive bacteria [44, 63], which may broaden the scope of therapeutic application. However, LspA inhibition does not universally sensitize Gram-positive bacteria to hGIIA killing, since hGIIA killing of *E. faecium* was not affected by myxovirescin A1 pretreatment. It is possible that myxovirescin could not reach LspA in sufficient amounts due differences in cell wall architecture between species and strains.

Alternatively, LspA has no role in hGIIA resistance in the *E. faecium* strain, therefore inhibition had no effect on susceptibility. Similar differences have been observed with regard to daptomycin resistance mechanisms, where mutations in the LiaFSR system caused a rearrangement of anionic membrane phospholipids in *E. faecalis* and daptomycin resistance but this was not observed for *E. faecium* [81]. More research is needed to clarify the potential application of LspA inhibitors as therapeutic add on for different Gram-positive pathogens.

hGIIA is considered as an acute phase protein [82]. It is strongly expressed by innate immune cells upon infection [10] and rises high levels in blood and organs that could be exploited for the development of new treatment strategies for MRSA infections. Deletion of *lspA* or its pharmacological inhibition renders MRSA more susceptible to hGIIA-mediated killing possibly due to altered membrane properties. Moreover, hGIIA resistance mechanisms overlap partially with daptomycin resistance mechanisms and indeed interference with LspA enhanced MRSA susceptibility to daptomycin. We only focused on hGIIA and clinically-relevant antibiotics, but it is possible that LspA inhibition has broader effects on virulence. We provided proof-of-concept for this potential add-on therapy by demonstrating that the antibiotics globomycin and myxovirescin A1 sensitizes MRSA for hGIIA-mediated killing, although strain-specific effects should be investigated. In addition to MRSA, *S. mutans* and *E. faecalis* were sensitized by pharmacological inhibition of LspA, increasing the impact of LspA as an sensitizing target. Therefore, interference with lipoprotein maturation through LspA inhibition is a strategy that warrants further exploration.

## Statements

### Author contributions

M.M.K., Y.W., V.P.v.H., G.S., C.P., and J.H. carried out the experiments. Y.W., C.P., G.L., J.H., R.M., and L.T. provided essential reagents. M.K. and V.H. took the lead in writing the manuscript. Y.W., G.L., J.H., R.M., Y.P., and L.T. revised the manuscript. N.S conceptualized the study and acquired funding. N.M.v.S., Y.P., and J.A.G.v.S. supervised the project.

### Data availability statement

Data and resources are available upon request from the corresponding author.

## Supplemental Figures

**Supplementary Table 1.**
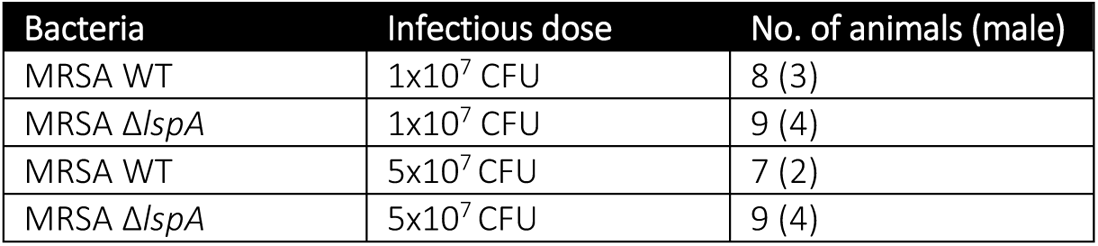
The four different groups including number and sex of mice used in the MRSA infection experiment using hGIIA-Tg C57BL/6 mice.

**Supplementary Table 2.**
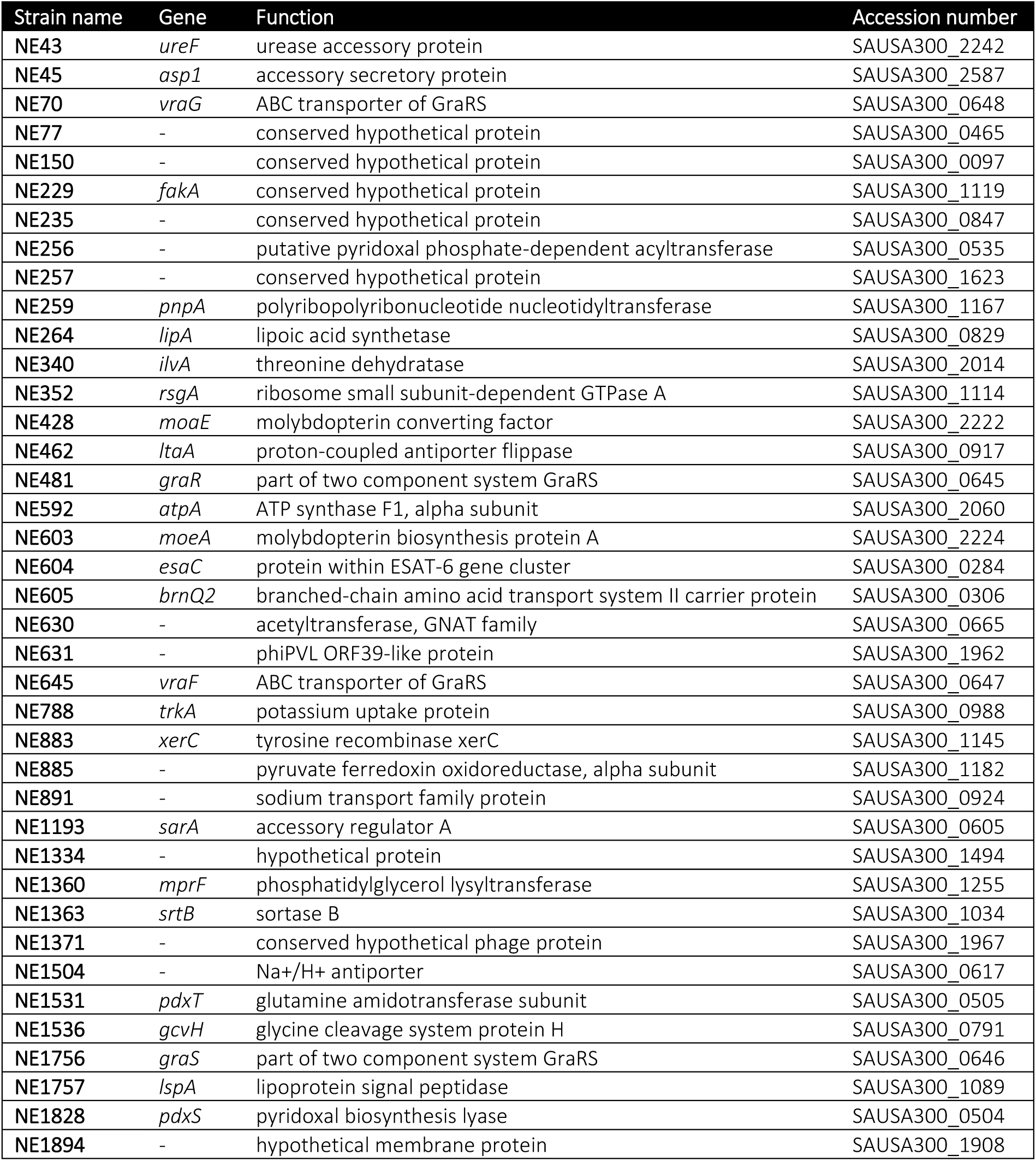
The 39 MRSA transposon mutants that showed decreased survival on plate after hGIIA exposure in a non-biased genetic screen.

**Supplementary Figure 1.**
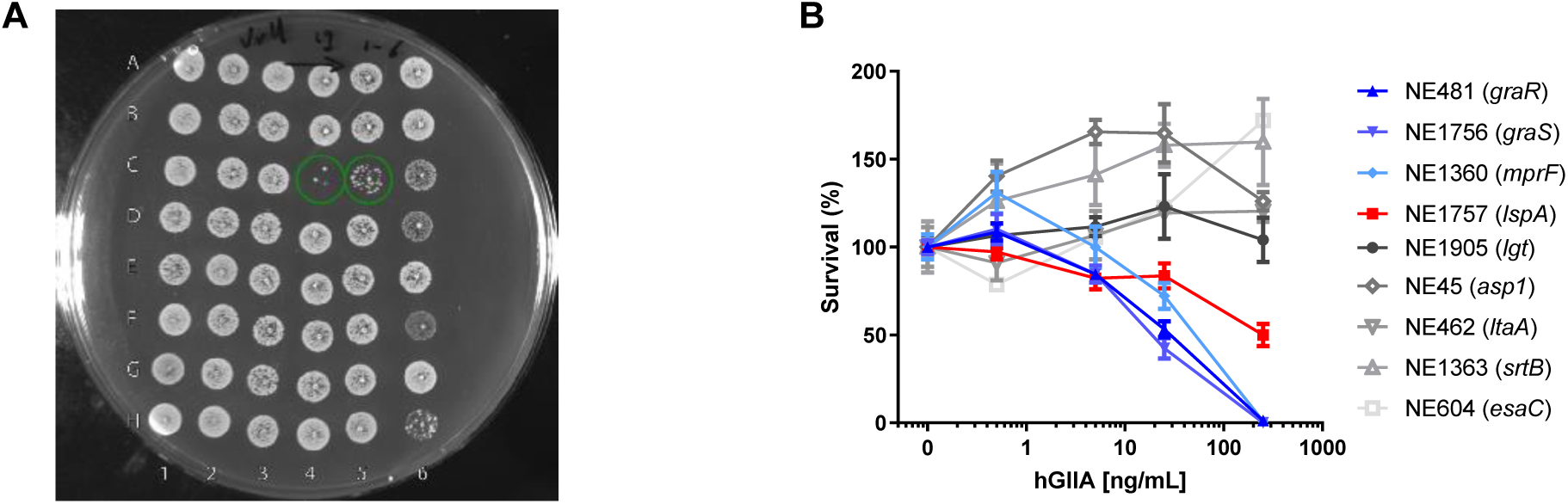
Identification of MRSA transposon mutants with increased susceptibility to hGIIA-mediated killing. (A) Representative image of an agar plate with spotted MRSA transposon mutants after exposure to 1.25 μg/mL recombinant hGIIA. Two transposon mutants on this plate showed decreased viability; at position C4 is NE0646 (*graS*) and at position C5 is the transposon mutant NE1757 (*lspA*). (B) Survival of potentially hGIIA susceptible mutants from the NTML using quantitative concentration-dependent killing assay. Data represent mean with standard error of the mean from three technical replicates.

**Supplementary Figure 2.**
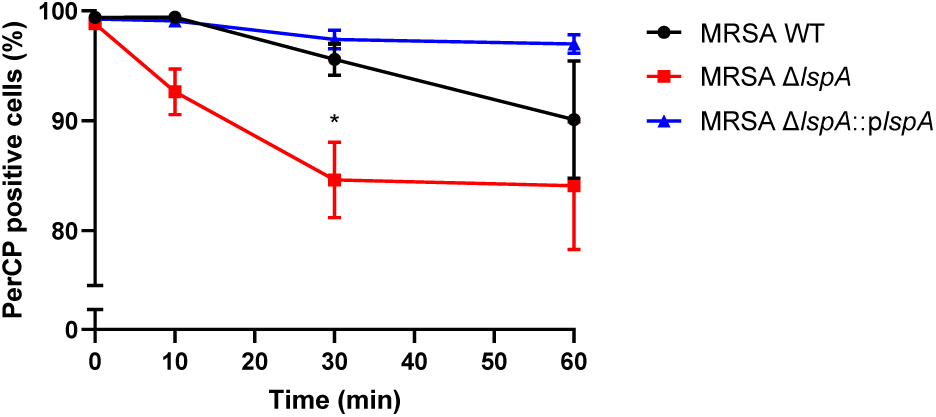
Faster membrane depolarization in *lspA* deletion mutant compared to WT and complemented strain after hGIIA exposure. Flow cytometric analysis of PerCP-positive cells of MRSA WT, MRSA Δ*lspA*, and MRSA Δ*lspA*::p*lspA* stained with DiOC_2_(3) at different time point after exposure to 500 ng/mL hGIIA. Data represent mean ± standard error of the mean of three independent experiments. Statistical significance was determined using a two-way ANOVA + Bonferroni’s Multiple Comparison Test. \**p* < 0.05. Data represent mean with standard error of the mean of three biological replicates.

**Supplementary Figure 3.**
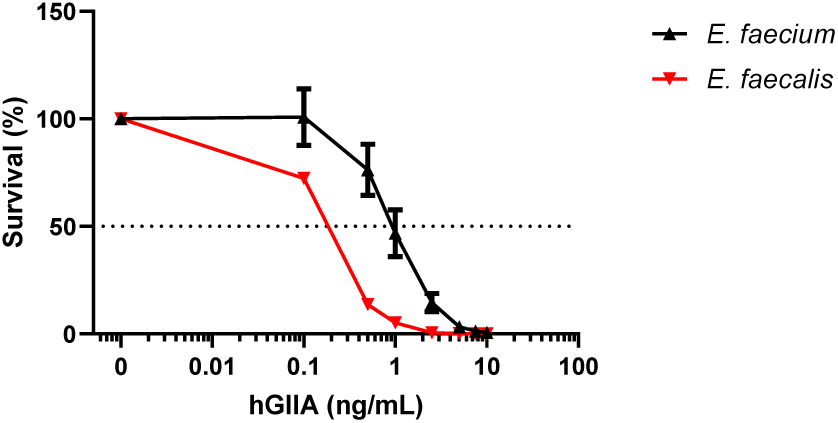
*E. faecalis* is more sensitive to hGIIA killing compared to *E. faecium*. Survival of *E. faecalis* V583 and *E. faecium* U0317 after exposure to concentration range of recombinant hGIIA. Data represent mean with standard error of the mean of three biological replicates.

